# Managing congestion at visitor hotspots using park-level use level data: Case study of a Chinese World Heritage site

**DOI:** 10.1101/596031

**Authors:** Jin-Hui Guo, Tian Guo, Kai-Miao Lin, Yu-Fai Leung, Qiu-Hua Chen

## Abstract

Tourist congestion at hot spots has been a major management concern for UNESCO World Heritage Sites and other iconic protected areas. A growing number of heritage sites employ technologies, such as cameras and electronic ticket-checking systems, to monitor user levels, but data collected by these monitoring technologies are often under-utilize. In this study, we illustrated how to integrate data from hot spots by camera-captured monitoring and entrance counts to manage use levels at a World Heritage Site in southeastern China. 6,930 photos of a congestion hotspot (scenic outlook on a trail) were collected within the park at a 10-minute interval over 105 days from January to November 2017. The entrance counts were used to predict daily average and maximum use level at the hotspot Average use level at the congestion hotspot did not exceed the use limit mandated by the Chinese park administration agency. However, from 9:20 am to 12:00 pm, the use level at hotspots exceeded visitor preferred use level. Visitor use level was significantly higher at the hotspot during a major Chinese “golden week” holiday. The daily entrance counts significantly predicted the average and maximum use level at the hotspot. Based on our findings, we recommend that the number of visitors entering the gate on each day should be less than 28,764 for the hotspots to meet use level mandates, while less than 6,245 to meet visitor preference. The gap manifested the complexity in visitor capacity management at high-use World Heritage Sites and other protected areas and calls for innovative monitoring and management strategies.

## Introduction

UNESCO World Heritage sites and similar iconic protected areas attract millions of visitors each year. While visitation generates economic and social benefits, the popularity invites issues such as overcrowding, congestion, and resources degradation. Particularly, when an excessive number of visitors congregate at “hotspots”, it can result into intensified environmental impacts, safety hazards, and dissatisfied users [1-3]. Since significant resources, such as magnificent views or iconic geological features, are typically near these hotspots, their overuse and potential spillover effects on the adjacent resource areas is a cause of concern. In China, there are 53 World Heritage Sites and more than 5000 other types of parks and visitor attractions which attracted billions of visitors annually, resulting in severe concerns of congestion and safety hazards related to tramping and falling [4-7]. Indeed, avoiding too many visitors staying at these locations at the same time has been a common management challenging many World heritage sites, especially in China with such a large population.

The congestion in park settings can seemed as too many people or vehicles gathering at a crowded place, while crowding highlights whether there are too many people by the personal judgment, which is a negative and subjective evaluation of a use level based on visitor perceptions [8-10]. Different approaches have been used to mitigate crowding and congestion at hotspots, such as educating visitors to avoid peak time, locations and improving infrastructure to increase capacity and managing the use level when use limits were set at park level [11]. In china, all parks and protected areas are mandated to devise a use limit developed by The National Guideline for Establishing Maximum Carrying Capacity for Parks (NGEMCC) which requires that each visitor should be able to occupy at least 0.5-1 m^2^ on the trail [12-14]. Within this range, park managers may devise their carrying capacity limits based on the park’s unique physical, social and biological characteristics. The priority of the guideline is to ensure visitor safety and alleviate congestion[14]. Many managers and researchers have noted that regulating visitor access on public lands is a contentious issue, but they didn’t address congestion issues at hotspots in parks base on this guideline[15-17]. As a matter of the fact, to gain public support for a use limits proposal [12], managers need to establish an empirical relationship that predicts how congestion and crowding level at hotspots are responsive to different use limits applied. However, the lack of empirical evidence to describe the relationship between use level and crowding is a vexing problem for managing visitors [18].

We acknowledge that the number of visitors does not equal with crowding severity. As an advancement in visitor management research, many researchers conceptualize crowding as a visitor subjective evaluation. Adopting this conceptualization helps managers to clarify the end goal for managing use level. However, empirical studies have also found little statistical relationships between visitor use level and the experience quality. An explanation is that there are many other factors, such as individual place attachments, activity types, and personalities, contributing to the perception of crowding[14, 19]. In practice, using proxy evaluative indicator, such as the number of people at one time(PAOT) at the hotspot or persons per viewscape (PPV) on trails, allow managers to fast evaluating the conditions and respond to potential issues of crowding [15]. With the multi-scale use level data, park managers could establish a link between visitation at park scale and at site scale to guide their long-term and short-term management decisions. However seldom evidence was found to address the crowding by monitoring total use level in western countries due to controversies about public privacy.

Different kinds of technology were applied to address the park visitor use management. The automated traffic counter was applied to examined the total number of visitors in United Stated Forest Service camp grounds at field personnel [20], it also was used to analyze the number of visitor use along the Carriage Roads in Acadia National Park [15]. Infrared technology on multiple-use trails was applied to reflect variability in the proportion of exiting day hikers across time of the day and the trail of use within Grand Canyon National Park [21]. More recently, the real-time and image-processing monitoring system was applied to underground station streams [22], urban communications [23] to estimate people flow and density. Using video to observe use level have compared counting accuracy with other methods [16]. However, they were seldom adopted to address crowding problem in most natural scenic spots, especially in eastern Asia to date. In china an amount of high-resolution cameras were installed at park by park managers and public security bureaus to acquire the visitor congestion distribution characteristics at multiple hot spots. High-resolution 24/7 cameras provide opportunities to expand and upgrade the existing visitor monitoring toolkits. Park managers who battled with visitor congestion at multiple hot spots on a daily basis could use the technologies to obtain visitor use data for the entire park and at specific sites. However, similar research is rare in eastern Asian countries, such as China with high population density and large visitation volume at parks and protected areas.

The objectives of this study are to manage congestion at visitor hotspots using park-level use level data acquired by high-resolution 24/7 cameras at World Heritage sites. Specifically, 1) to identify the empirical relationship between total use level and use level at hotspots in a World Heritage Site in southeastern China, and 2) to derive recommended use limits based on the identified relationship.

## Methods

### Park Management in China

Mount Wuyi National Park (27°32’36″∼27°55′15″N, 117°24′12″∼118°02′50″E) is in southeast of China (Fig. 1), approximately covering about 990 km^2^. It was designated as a World Natural and Cultural Heritage Site in 1999 for its outstanding biodiversity value, scenic beauty and cultural significance. Being close to major population centers along the east coast of China, the park faces constant pressure from high use volume and congestion on popular trails.

**Fig. 1.**
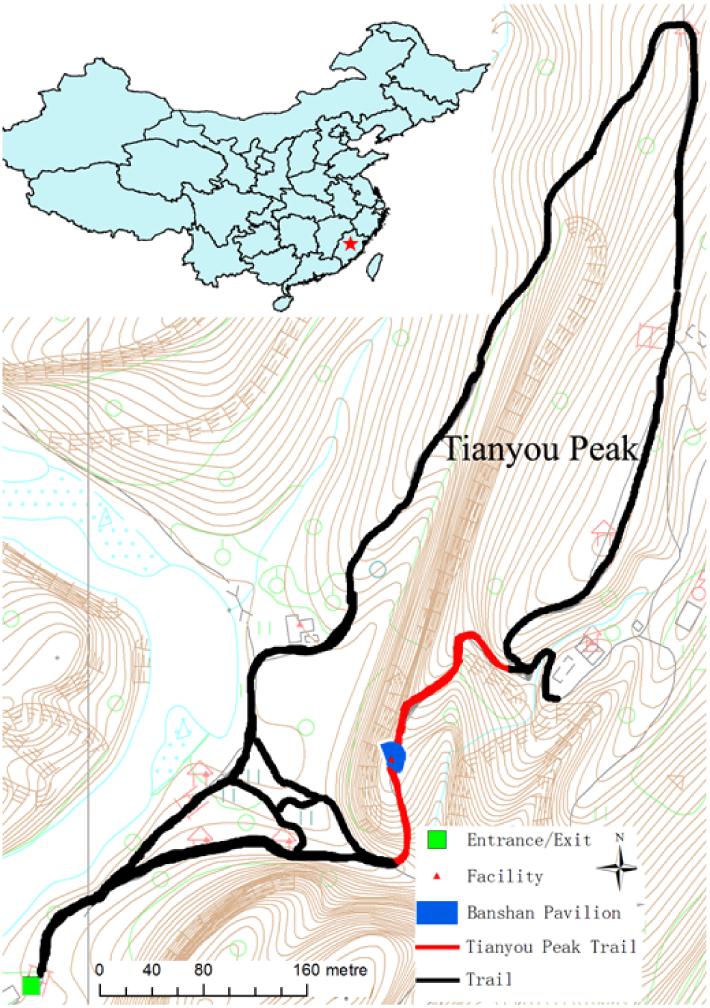
Trail condition and location of Tianyou Peak at Mount Wuyi National Park, China

The park has a ticket-checking system that monitors the number of visitors who have entered the park. Managers can impose desired use level limits through controlling how many tickets they sell. The ticket-checking systems at entrances help keep visitation records. High-resolution cameras at specific sites feed real-time images to managers, allowing managers to respond quickly if they observe a safety hazard. Together, the ticket-checking systems and cameras could obtain use-level data for the entire parks and specific sites.

Tianyou Peak Trail with 1.5-m-wide stone steps ascending 520 meters to the peak is one of the most visited attractions within the park. The outlook Banshan Pavilion rising about 200-meters from the trailhead is the most congested hotspot along the trail (Fig. 1 and Fig. 2). Many visitors get use to stop at this outlook to rest, enjoy the view and take photos. Thus, visitor congestion occurs often along the trail within the peak time during the holidays according to the monitoring videos from the high-resolution cameras. Besides high daily usage on weekends, managers estimated that over 20,000 tourists visited the trail every day during the weeklong Chinese New Year Holidays and Chinese National Day Holidays. These two important holidays, commonly referred to as the “golden weeks”, are critical times for generating tourism revenue every year. Concerned with high use volume and congestion on the trail, the park managers installed a high-resolution camera on the top of Tianyou Peak in 2016. The camera can rotate or be set manually and take pictures of multiple sites surrounding the Peak during park operation time.

**Fig. 2.**
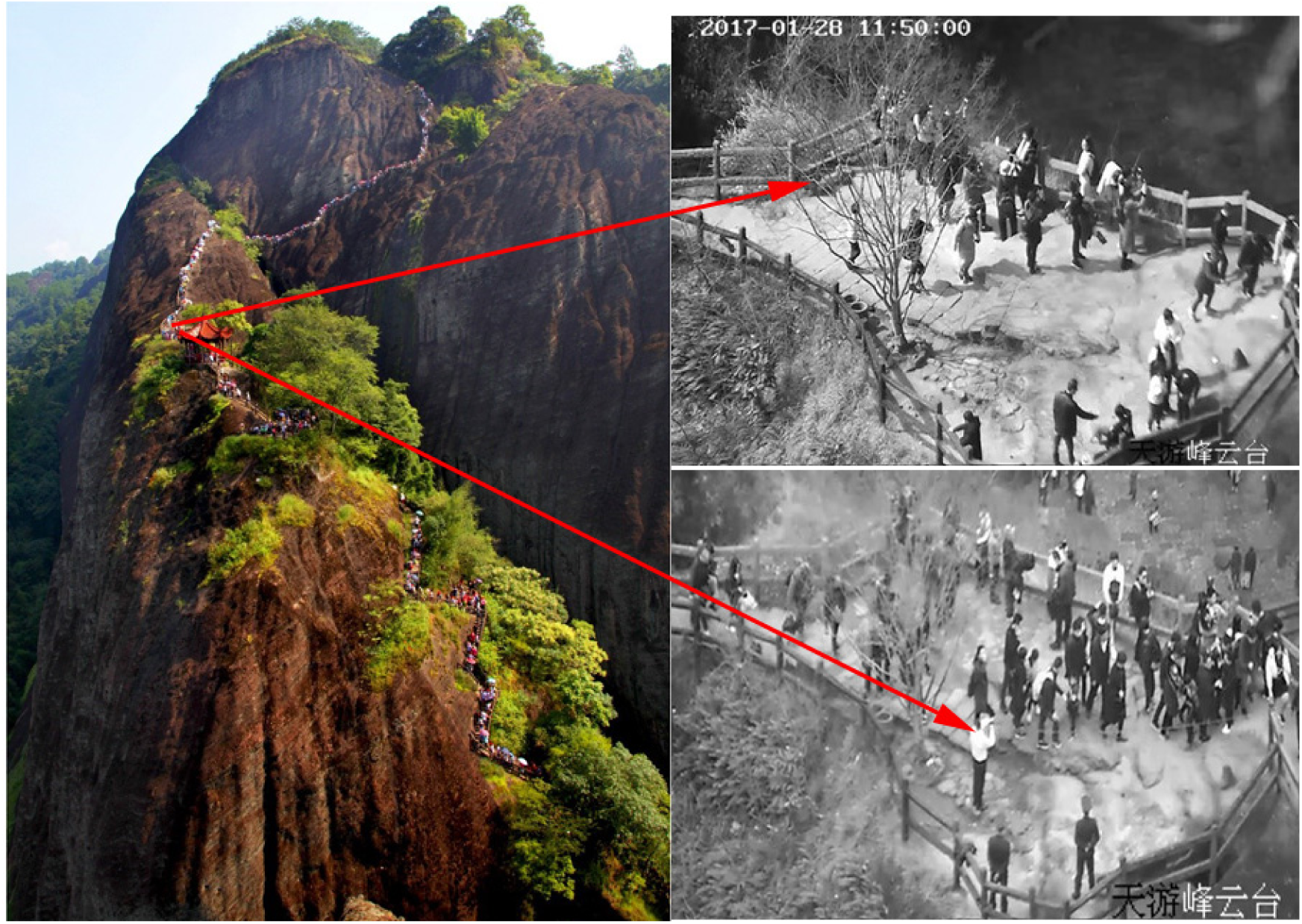
Illustration of photos of Banshan Pavilion on Tianyou Peak Trail in Mt. Wuyi National Park.

### Data Collection

We collected use level data for the park and at the Banshan Pavilion outlook from the park’s camera system between January 28^th^ and November 10^th^ 2017. In order to make data more representative, we worked with the park staff to keep the camera pointing to the outlook for one day with three days intervals, with a total of about 10-12 days per month. Consequently, photos were available for a total of 105 days. The photos were captured by surveillance at 10 min intervals from 7:00 am to 5:00 pm, producing 61 photographs per day to estimate average use level at the outlook throughout a whole day. A total of 6385 photographs were collected in all 105 days with a few photos missing due to equipment maintenance and occasional malfunctioning (Figure. 2). Eight research assistants worked in pairs to count the number of visitors in each photo, with one assistant counting and the checking accuracy. The photos captured 73,158 visitors in total. In addition, entrance counts data were provided through ticket-checking at park entrance for the same days.

In order to establish the acceptable and preferred standards of crowding, visual-based questionnaire was designed including the acceptability of different level of crowding and the respondents’ profiles. The questionnaire survey was administered for 5 weekends from November 5th to December 18th in 2016. 465 Visitors were surveyed on the site when they are visiting the summit of the Peak, which is about 30minutes from the outlook. Respondents were shown the photos with different visitor use level of the trail slope. They were asked to rate the acceptability from the most unacceptable (−4) to the most acceptable (+4) and the neutral point of 0. In addition, we measured the area of the Banshan Pavilion in order to evaluate the limiting standard of crowding.

### Data Analysis

We first conducted descriptive analysis on the use level of the outlook leading to Tianyou Peak. We calculated visitor use level by averaging people per photo and visualized the visitor use level at the outlook over time (Table 1). We then compared the average visitor use level against three crowding standards, that is, limiting standard, the acceptable standard and the preferred standard. We did a separate analysis on the temporal use level distribution for data collected during the Chinese New Year Holidays from January 28^th^ to February 3^rd^, 2017 and China National Day Holidays from October 1^st^ to October 7^th,^ 2017.

We conducted linear regression statistical analysis in order to link the use level of the entire park with that of the outlook. Since the entrance counts were only available on the daily level, we aggregated the people per photo data to the daily level. Specifically, we summed up people per photo collected in the same day and divided it by the number of photos collected in that day, obtaining the average number of visitors at the outlook at any given time in a day (hereafter *average use level)*. We also obtained the maximum number of visitors at the outlook at a given time in a day (*hereafter maximum use level)*. We fit two models with the daily entrance counts as independent variable, and average use level and maximum use level as dependent variables respectively.

## Results

### Temporal visitor use level

Questionnaire survey results showed that park visitors to Tianyou Peak preferred less than 27 people at the outlook (*hereafter preferred standard*), while they considered less than 45 people at the outlook acceptable (*hereafter acceptable standard*). The 14-meter-long and 4-meter-wide Banshan Pavilion covers 56 m^2^ through field observation. We evaluated that each visitor should occupy at least 1 m^2^ at the outlook considering the Situations based on NGEMCC. We calculated that the maximum number of visitors at the outlook at any given moment should not exceed 56 people for the safety (*hereafter limiting standard*).

The daily temporal visitor use level at the outlook against three crowding standards were showed at Fig.3. In normal times, the average use level begins to grow at 8:00 am and a peak comes out at 10:00 am, which never exceeds the limiting standard and rarely exceeds the visitor acceptable standard but exceeds the visitor preferred standard. The visitor use level keeps nearly at a constant from 12:00 pm to 15:30 pm and then falls quickly till the end. During this period the average use level never goes beyond the visitor preferred standard. However, during the Chinese New Year Holidays, visitor use level at the outlook peaked at 10:50am which exceeds the limiting standard by 178% and exceeds all three standards nearly from 8:40 am to 14:40 pm. During Chinese National Day Holidays, the use level peaked at 11:20am which exceeds the limiting standard by 64% and goes beyond all limiting standards from about 9:20 am to 12:00 pm (Figure 3).

**Fig. 3.**
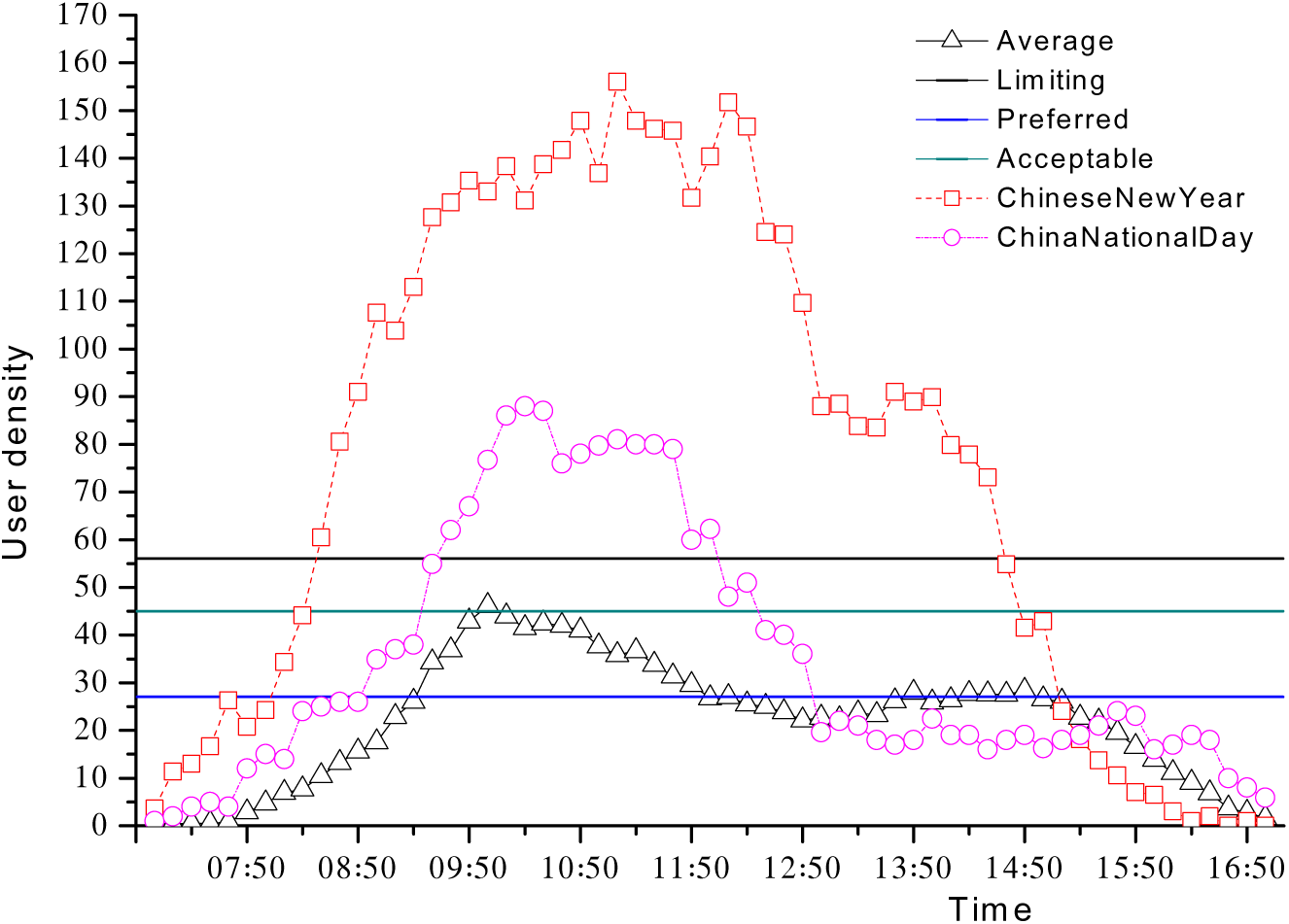
Use level at Banshan Pavilion on the Tianyou Peak Trail at Mt. Wuyi National Park, China. Average: Daily average visitor use level in normal times, preferred: Visitor preferred standard; Acceptable: Visitor acceptable standard; Limiting: Limiting standard; Chinese New Year: Visitor use level over the Chinese New Year Holidays; China National Day: Visitor use level over the National Day Holidays.

### Correlation between use level for the entire park and at the trail

The linear regression analysis revealed a significant relationship between the use level for the entire park and at the outlook. The entrance counts significantly predicted daily average visitor use level of the outlook (*F* (1,105) =288.604, *p-value*<.000, *R*^*2*^=0.829). For every additional 1000 visitors passing through the entrance, on average there will be about 5 more visitors at the outlook at a given time (Fig. 4). The entrance counts also significantly predicted the maximum use level at the outlook (*F* (1,105) =412.57, *p-value*<.000, *R*^*2*^=0.798). For every additional 1000 visitors passing through the entrance, there will be 6 more visitors at the outlook when the uses of the hotspot peaks (Fig. 4).

**Fig. 4.**
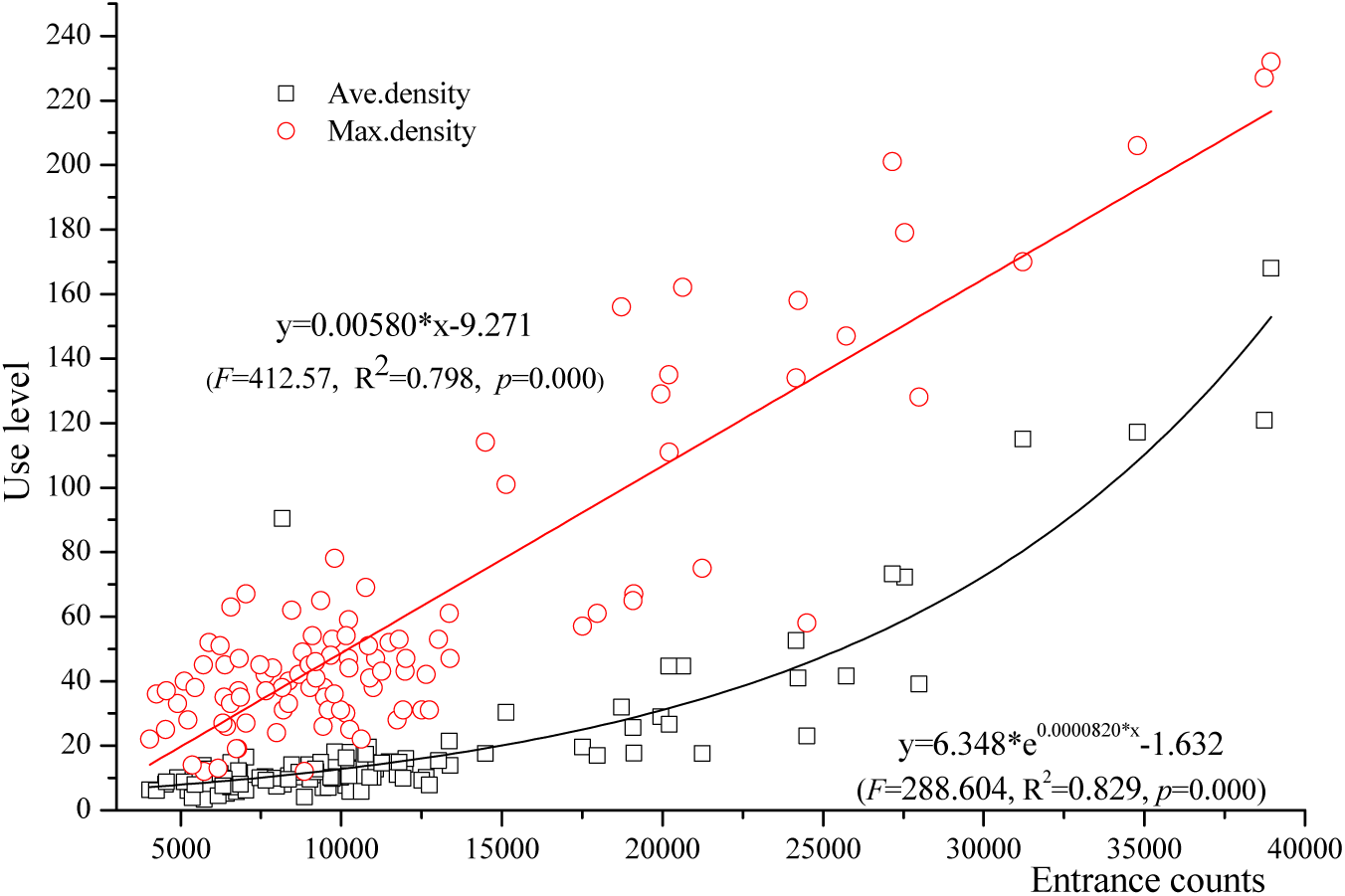
Model results of using entrance counts to predict average use level and maximum use level at the outlook

### Classification of Congestion degree

Using the regression model, we calculated at most how many visitors passing by the entrance daily should be permitted so that the average and maximum use level at the outlook will not exceed the crowding standards (Table 3).

**Table 3.**
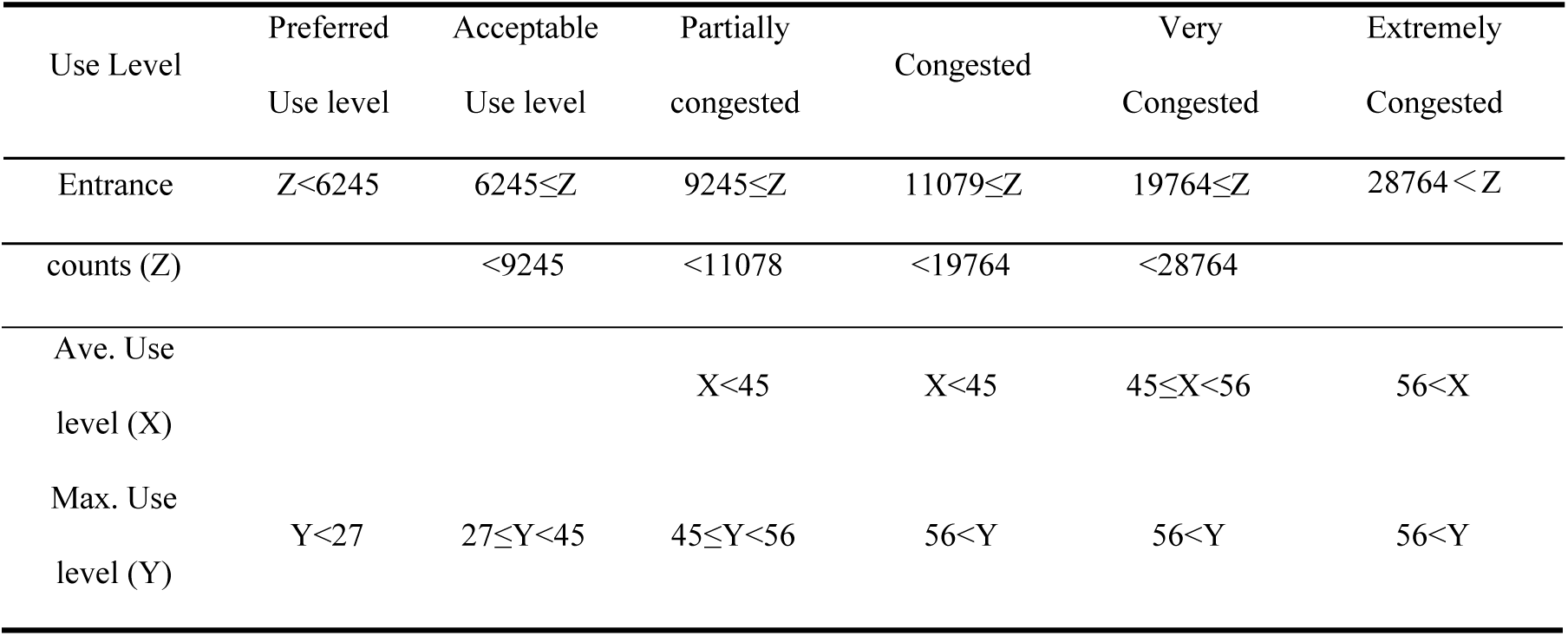
Threshold values of entrance counts for combinations of average use level and maximum use level values at the trail

1) *Preferred*, when the number of visitors passing by the entrance is less than 6,245, the maximum use level and the average use level at Banshan Pavilion meet the visitor preferred standard. It means congestion never occurs during a whole day.

2) *Acceptable*, when the daily number of visitors passing by the entrance is between 6,245 and 9,245, the maximum use level at the outlook will exceed the visitor preferred standard, but meet the visitor acceptable standard. It means congestion may occurs for a short time but visitors feel acceptable.

3) *Partially Congested*, when the daily number of visitors passing by the entrance is between 9,245 and 11,078, the maximum use level at the outlook will exceed the visitor acceptable standard, but meets the limiting standard. It means congestion only occurs during peak hours but safety management can be guaranteed with any management measures.

4) *Congested*, when the daily number of visitors passing by the entrance is between 11,078 and 19,764, the maximum use level at the outlook will exceed the limiting standard while the average use level meets the visitor acceptable standard. It means congestion takes place during peak time, but on average visitors can accept the occasion.

5) *Very Congested*, when the daily number of visitors passing by the entrance is between 19,764 and 28,764, the maximum use level at the outlook will exceed the limiting standard but the average use level meets the limiting standard. That means extreme congestion occurs, but on average visitors’ safety can be guaranteed if management measures are taken action.

6) *Extremely Congested*, when the daily number of visitors passing by the entrance is over 28, 764, the average and maximum use level at Banshan Pavilion will both exceed the limiting standard. It means whatever action was taken, congestion can not be alleviated affectively.

## Discussion

### Temporal visitor use level at the hotspot

Results based on the camera-captured monitoring data showed that the temporal visitor use level between normal times and two golden week holidays both presented the characteristics of the single peak distribution. The use level concentrated during 9:00am - 12:30pm. An explanation is that visitors preferred to climb the trail to enjoy the Danxia landscape scenery in the early morning. Another reason is due to the special landform as the Peak is composed of a great rock without tree shade along the trail. Thus, visitors prefer to climb the trail avoiding exposure under the sunshine.

However, results indicated that there is a distinct-difference about the temporal visitor use level between normal times and two golden week holidays. The visitor crowding was very high in two golden week holidays because of the influx of visitors. On one hand, Park managers can not adopt the proactive management decision-making because of lacking the quantitative relationship between the entrance counts and the crowding level at the hotspot. On the other hand, visitor swarming into the park entrance were not well informed about the congestion level at the hotpots for the lack of necessary information. Therefore, the quantitative relationship between the use level of the entire park and that of the hotspot was urgently established to predict the crowding level at the hotspot during peak seasons.

### Links between the use level of the entire park and that of the hotspot

Quantitative information on the hotspot and intensity of visitor use can alert park administers to manage areas prone to congestion, crowding or safety concerns [19]. Statistical modeling using regression analysis is an effective use of visitor use monitoring data. Results suggested that the relationship between the use level of the entire park and that at the hotspot, once established, can predict congestion level of a tourist hotspot and evaluate compliance or violation of pre-determined use limits based on different management objectives. Only a few researches have tried to predict the visitor use level at the hotspots in western countries [15, 24-26]. The study indicated that establishing links between the entire park and that of the hotspot is feasible in China with large population context. Furthermore, most of parks in China charge fees by checking tickets at the entrances which make it easy to control the total use level to achieve the anticipated management objectives. Therefore, the findings provided Chinese park managers with information to help alleviate congestion, reduce crowding and improve safety conditions at the hotspot under park-scale use limits.

### Visitor capacity management at the hotpot

Visitor capacity research is growing in China [27-30]. However, most of these studies focused on park-scale use level, lacking attention to site-scale use level. The study helped to fill in this gap. This paper compared the visitor use level and the threshold values of entrance count and the results showed that the daily entrance counts significantly predicted the average use level and maximum use level at the hotspot. The use level at the outlook exceeded limiting standard for most of the day during the Chinese New Year Holiday. For most time of an average day, use level at the hotspot did not exceed limiting standard, visitor acceptable standard, or the visitor preferred standard, except for a short period when the use level at the hotspot exceeded the visitor preferred standard. Therefore, less than 6,245 to meet the visitor preferred standard, while the number of visitors permitted each day should be less than 28,764 for the visitor safety. If visitor use level exceeded the thresholds, the park management must be faced with the great challenge in countering safety and mitigating impacts of overcrowding.

### Management Implications

This study demonstrated how managers could couple camera-captured monitoring data with entrance counts to manage visitor capacity at World Heritage Sites and parks with high use volume. Different congestion scenarios will require different management strategies. For example, in the *Partially Congested* scenario, the maximum number of visitors at the congestion hotspot will exceed visitor acceptable standard, but the average number of visitors at the hotspot will not exceed the acceptable standard. Managers could focus on helping visitors to avoid anticipated peak hours, using behavioral change approaches such as time-based ticket discount for those who visit the park early in the morning. Information may also help visitors avoid hours prone to congestion, such as using a mobile app to distribute real time images of hotspots. In the *Extremely Congested* scenario, park managers should prepare the fast-response protocol.

Camera could help park managers at other World Heritage Sites to monitor use level at congestion hotspot. Limited park budgets, policies and regulations, or unique size and shape of the congestion hotspots may prevent managers to employ cameras to monitor use level. For example, processing visual data may be labor intensive. We had eight research assistants counting people per photo for approximately 346 hours. Crowdsourcing such as Amazon Mechanic Turk may be another way to complete the counting task [31]. Violating privacy may be a concern for some cultures. The camera was set up on the top of Tianyou Peak at Mt. Wuyi National Park. The long distance between the camera and the outlook makes it almost impossible to identify individuals in the photos. Managers at other parks who are interested in using cameras for visitor monitoring could carefully design the distance and angle of the camera to avoid invading individual privacy.

### Limitations and Future Studies

This study contributed to the understanding of visitor capacity at highly populated Asian countries through focusing on visitor monitoring and the number of people at one time at attraction sites. We did not examine other important visitor experience indicators such as the amount of recreational impacts along the trail or maximum waiting time on the trail [32]. We acknowledge that visitor monitoring using camera and entrance counts should complement other types of planning and monitoring strategies in visitor capacity management. We did not randomly select days to collect camera-captured data. There were no reasons to suspect we introduced bias to our data through sampling. The availability of entrance counts limited our regression analysis. We aggregated people per photo to variables at the daily level – the average visitor use level and maximum visitor use level. Future studies could improve the regression model through using a finer time scale, such as linking entrance counts per hour with the number of visitors at any given time at attraction sties. The results supported a correlation between park use level and site use level, but it did not explain the mechanism behind this link. Future research using GPS and agent-based modeling could help identify visitor movement pattern and predict when congestion occurs on the trail.

## Conclusions

UNESCO World Heritage Sites and other iconic protected areas face the constant tension between visitor use and protecting environment and visitor experiences. Monitoring use level is a core management strategy for visitor carrying capacity. We found camera-captured data is useful to monitor use level at a congestion hotspot within a highly visited World Heritage Site in southeast China. The average and maximum use level at the hotspot could be predicted by entrance counts, suggesting direct relationships between park-scale use level and site-scale use level. Our results suggested park managers should not rely on a single number to manage congestion at hotspots. Instead, park managers need a broader decision-making process including multi-scale visitor monitoring. We need more research to advance our understanding of visitor capacity at different cultural and management contexts and help World Heritage Sites to address the congestion problem.

## Uncategorized References

1. Ryan C, Cessford G. Developing a visitor satisfaction monitoring methodology: quality gaps, crowding and some results. Current Issues in Tourism. 2003;6(6):457–507. https://doi.org/10.1080/13683500308667966.

2. Manning R, Wang B, Valliere W, Lawson S, Newman P. Research to estimate and manage carrying capacity of a tourist attraction: A study of alcatraz Island. Journal of Sustainable Tourism. 2002;10(5):388–404. https://doi.org/10.1080/09669580208667175.

3. Kohlhardt R, Honeyrosés J, Lozada SF, Haider W, Stevens M. Is this trail too crowded? A choice experiment to evaluate tradeoffs and preferences of park visitors in Garibaldi Park, British Columbia. Journal of Environmental Planning & Management. 2018;61:1–24. https://doi.org/10.1080/09640568.2017.1284047.

4. Zhang B, Wu H, Fei X, Mo S. Protecting the national parks and the world heritage sites in China: challenge and strategy. Journal of Geographical Sciences. 2004;14(S1):87–90. https://doi.org/10.1007/bf02841113.

5. Cros HD, Leask A, Fyall A. Too much of a good thing? Visitor congestion management issues for popular World Heritage tourist attractions. Journal of Heritage Tourism. 2008;2(3):225–38. https://doi.org/10.2167/jht062.0.

6. Li M, Wu B, Cai L. Tourism development of World Heritage Sites in China: A geographic perspective. Tourism Management. 2008;29(2):308–19. https://doi.org/10.1016/j.tourman.2007.03.013.

7. Agnew N, Demas M. Visitor management and carrying capacity at World Heritage Sites in China: Extended abstracts of the international colloquium. The Getty Conservation Institute; Los Angeles, CA2014.

8. Lime DW. Congestion and crowding in the National park system. University of Minnesota Digital Conservancy: University of Minnesota. Agricultural Experiment Station, 1996.

9. U.S. National Park Service. Congestion management toolkit U.S. National Park Service website: https://www.nps.gov/transportation/pdfs/NPS-CMS_Toolkit.pdf: U.S. National Park Service 2014.

10. World Tourism Organization. Tourism congestion management at natural and cultural sites: a guidebook. Tourism Congestion Management at Natural & Cultural Sites A Guidebook. 2004.

11. Manning RE. Defining and managing visitor capacity in National Parks: A program of research in the U.S. National Park System. 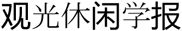. 2011;17(2):183–214. https://doi.org/10.6267/jtls.2011.17(2)4.

12. Guo Nei, Lv You, Kuang Q. Domestic tourism statistics China National Bureau of Statistics’ website: http://data.stats.gov.cn/easyquery.htm?cn=C01: China National Bureau of Statistics; 2016.

13. Tian K, Tian J. Study on environment capacity of core attractions of Wulingyuan Scenic and Historical Interest Area. Journal of Green Science & Technology. 2011.

14. Cui F, Yang Y. A study on the time-space distribution features and utility intensity of the TEBC Resource of Mt. Tai (in Chinese). Geographical Research. 1997;16(4):47–55. https://doi.org/10.11821/yj1997040007.

15. Pettebone D, Meldrum B, Leslie C, Lawson SR, Newman P, Reigner N, et al. A visitor use monitoring approach on the Half Dome cables to reduce crowding and inform park planning decisions in Yosemite National Park. Landscape & Urban Planning. 2013;118(118):1–9. https://doi.org/10.1016/j.landurbplan.2013.05.001.

16. Arnberger A, Haider W. A comparison of global and actual measures of perceived crowding of urban forest visitors. Journal of Leisure Research. 2007;39(4):668–85. https://doi.org/10.1080/00222216.2007.11950127.

17. Beunen R, Regnerus HD, Jaarsma CF. Gateways as a means of visitor management in national parks and protected areas. Tourism Management. 2008;29(1):138–45. https://doi.org/10.1016/j.tourman.2007.03.017.

18. Sim K-W, Kim B-G, Lee J-H, Poung-Sik Y. The evaluation of effectiveness of the interpretive program at national parks. Journal of Outdoor Recreation and Tourism. 2018;21:69–75. https://doi.org/10.1016/j.jort.2018.01.004.

19. Schultz J, Svajda J. Examining crowding among winter recreationists in Rocky Mountain National Park. Tourism Recreation Research. 2017;42(1):84–95. https://doi.org/10.1080/02508281.2016.1259029.

20. Tombaugh LW, Love LD. Estimating number of visitors to National Forest campgrounds. Research note RM 1964:17-9.

21. Schwartz Z, Stewart W, Backlund EA. Monitoring visitor flows in destinations: the case of multiple-use hiking trails in Grand Canyon National Park. Tourism Analysis. 2009;14(6):749–63. https://doi.org/10.3727/108354210x12645141401142.

22. Boninsegna AM, Coianiz T, Trentin E. Estimating the crowding level with a neuro-fuzzy classifier. Journal of Electronic Imaging. 1997;6(3):319–28. https://doi.org/10.1117/12.269900.

23. Yun Z, editor Information system for monitoring the urban environment based on satellite remote sensing: Shanghai as an example. Geoscience & Remote Sensing, Igarss 97 Remote Sensing-a Scientific Vision for Sustainable Development, IEEE International; 1997.

24. Broom TJ. An assessment of indirect measures for the social indicator of encounters in the Tuolumne Meadows area of Yosemite National Park: University of Idaho; 2010.

25. Pettebone D, Newman P, Lawson SR. Estimating visitor use at attraction sites and trailheads in Yosemite National Park using automated visitor counters. Landscape & Urban Planning. 2010;97(4):229–38.

26. Lawson S, Newman P, Choi J, Pettebone D, Meldrum B. Integrated transportation and user capacity research in Yosemite National Park: the numbers game No.2119. Transportation Research Record. 2009. https://doi.org/10.3141/2119-11.

27. Huang ZF, Yuan LW, Jun-Lian GE, Qiu-Shi GU. Tourism environmental carrying capacity and its assessment research——A case study of ecotourism in coastal wetland ofjiangsu. Scientia Geographica Sinica. 2008;28(4):578–84. https://doi.org/10.3969/j.issn.1000-0690.2008.04.021.

28. Lili, CUI, Lijuan, WU, Ming. Tourist behaviors in Wetland Park:A preliminary study in Xixi National Wetland Park, Hangzhou, China. Chinese Geographical Science. 2010;20(1):66–73. https://doi.org/10.1007/s11769-010-0066-4.

29. Pan LL, Yang-Mei MA. Psychological capacity of tourist in Xixi National Wetland Park based on crowding perception. Wetland Science. 2014;12(5):662–8. https://doi.org/10.13248/j.cnki.wetlandsci.2014.05.019.

30. Luo J, editor Simulation of congestion of visitors moving in 2010 Shanghai Expo. International Conference on E-Business and Information System Security; 2009.

31. Hipp JA, Adlakha D, Eyler AA, Gernes R, Kargol A, Stylianou AH, et al. Learning from outdoor webcams: Surveillance of physical activity across environments 2017.

32. Manning R, Valliere W, Minteer B, Wang B, Jacobi C. Crowding in parks and outdoor recreation: a theoretical, empirical, and managerial analysis. Journal of Park and Recreation Administration. 2000;18(4):57–72.

